# Age-dependent accumulation of neuromelanin in African mole-rats

**DOI:** 10.64898/2026.10.13.759023

**Authors:** C. Canales, Mario M. Ilarduya, J. Chocarro, A.J. Rico, G. Ariznabarreta, E. Lorenzo-Ramos, J. Hoyo, D. Carbonell, R. Sumbera, J. Okrouhlik, D. Ruiz-Molina, J.L. Lanciego

## Abstract

Although neuromelanin (NMel) is a dark pigment characteristic of catecholaminergic cell groups in the primate CNS, rodent brains lack pigmentation in ventral mesencephalic cell groups and in the locus coeruleus (LC). Even though these neurons possess the molecular machinery required to produce NMel, it remains unknown why the brains of mice and rats are not pigmented. Accordingly, postmortem brain samples of two species of African mole-rats (*Heterocephalus glaber* and *Fukomys mechowii*) were used to evaluate whether these longest-living rodents accumulate NMel. The results revealed time-dependent pigmentation beyond two years of age, with the highest pigmentation levels observed in LC neurons. In conclusion, the physiological pigmentation of catecholaminergic neurons in the brain of African mole-rats appears to result from a slow kinetic process that requires a lifespan long enough to allow detectable NMel accumulation.

## 1 Introduction

In the primate brain, catecholaminergic neurons of the substantia nigra pars compacta (SNpc; *nigra/nigrae* meaning ‘dark’ or ‘obscure’), the ventral tegmental area (VTA), and the locus coeruleus (LC; *coeruleus/ceruleus* meaning ‘dark blue’) display robust neuromelanin (NMel) pigmentation. NMel most likely forms through non-enzymatic auto-oxidation of cytosolic dopamine *(1,2)*. Moderate-to-high levels accumulate exclusively in primate brains (human and non-human) in a time-dependent manner *(3)*, with the first detectable traces of NMel observed in human postmortem samples older than three years *(4)*.

Work conducted in the past few decades has substantially advanced our understanding of the role of NMel in both healthy and diseased brain states. However, the complex biology underlying the pigmentation of catecholaminergic cell groups remains incompletely understood *(5)*. This gap is particularly relevant given the proposed direct link between NMel accumulation and dopaminergic neuronal vulnerability in postmortem human samples from patients with Parkinson’s disease (PD) *(6)*. These findings suggest that NMel is not merely a passive pigment or metabolic byproduct, but may instead be an active participant in the cascade leading to dopaminergic degeneration. If NMel is indeed causally associated with PD, a fundamental question arises: should NMel formation be regarded as a primate-specific adaptation, perhaps intrinsically linked to species that develop PD-like pathology, or as a broader, evolutionarily conserved biological process? Resolving this question would clarify the evolutionary significance of NMel and its relationship to catecholamine metabolism and PD. A paradigmatic case is that of rats and mice, which are generally thought not to produce NMel and, consequently, do not naturally develop progressive nigrostriatal dopaminergic neurodegeneration. Interest in this question has recently resurged with the development of rodent and non-human primate models of PD based on viral-mediated tyrosinase overexpression. In these models, forced NMel accumulation reproduces all the major neuropathological hallmarks of human PD with remarkable fidelity *(7-10)*. Thus, does the apparent absence of NMel in mice and rats and other short-lived species reflect a genuine inability to synthesize the pigment, or is it instead a consequence of the inherently slow nature of dopamine auto-oxidation, such that NMel becomes detectable only after years or decades of continuous catecholamine turnover?

To assess whether the short lifespan of mice and rats might explain their lack of brain pigmentation, postmortem samples from two species of African mole-rats (*Heterocephalus glaber* and *Fukomys mechowii*) were analyzed. Both mole-rats are highly social mammals that live in underground families of up to 300 individuals, in which a single female and a single male are typically the only reproducing members. In terms of longevity, the ratio of maximum observed lifespan to weight-predicted lifespan exceeds four, a value comparable to that of humans *(11)*. Mole-rats display a remarkable suite of physiological adaptations. *H. glaber* is highly resistant to cancer *(12-14)*, maintains vascular health throughout life *(15)*, it is well adapted to limited oxygen availability *(11)*, and can reduce its metabolic rate by up to 25% in response to prolonged periods of food deprivation *(16)*. They also exhibit remarkable resilience to age-associated neurodegenerative disorders such as Alzheimer’s disease. Although they accumulate high levels of amyloid-beta peptide and tau protein, they do not develop extracellular amyloid plaques or intracellular neurofibrillary tangles *(17,18)*, likely owing to robust mechanisms that mitigate hypoxia and oxidative stress *(17)*. These unique characteristics contribute to an exceptionally long lifespan, with individuals living on average between 25 and 30 years *(11,19)*. Although less extensively studied, many of these adaptations, including exceptional longevity, are shared by other mole-rat species such as *F. mechowii (20)*.

## 2 Materials and Methods

### Study design

This study investigates why mice and rats -the most commonly used laboratory model organisms-lack neuromelanin (NMel) pigmentation in their catecholaminergic brain centers. NMel is most likely a by-product of dopamine auto-oxidation; thus, although rodents possess the molecular machinery required for NMel synthesis, their brains remain unpigmented, for reasons that are not understood. Because NMel accumulates through a slow kinetic process, our working hypothesis is that mice and rats simply do not live long enough to develop detectable pigmentation. To this end, postmortem brain samples from two mole-rat species, *Heterocephalus glaber* and *Fukomys mechowii*, both renowned for their exceptionally long lifespan, were analyzed (Figure 1).

**Figure 1.**
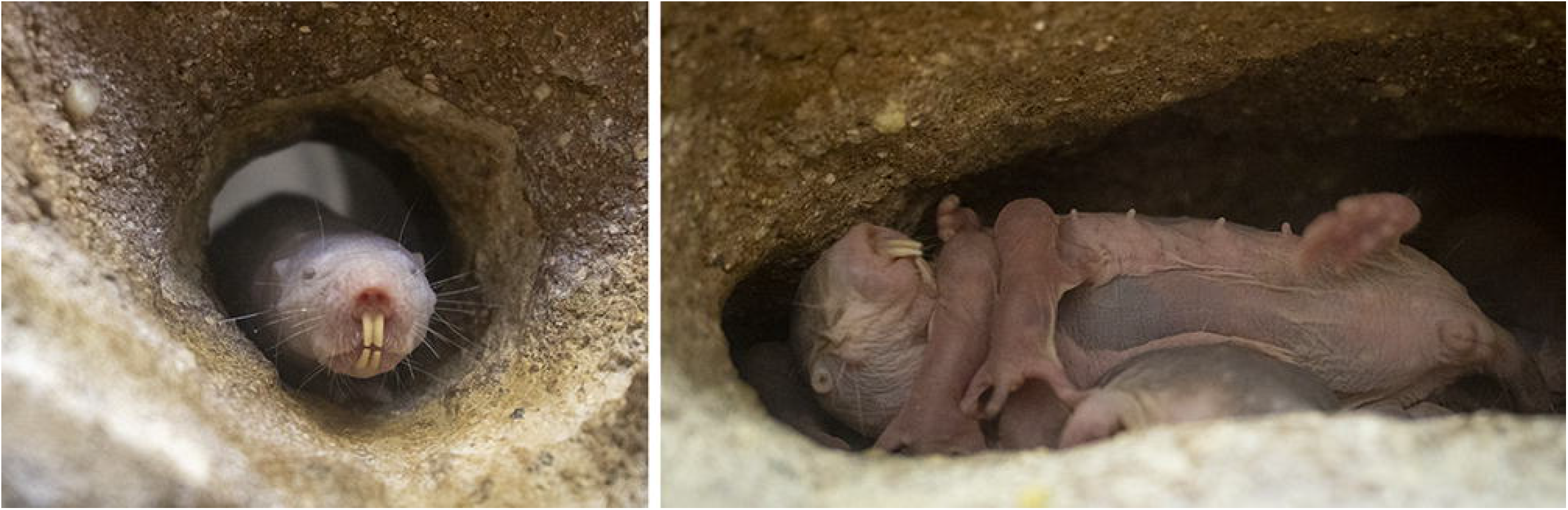
The naked-mole rat (*H. glaber*). Considered the world’s longest-living rodent, the naked mole-rat is a small subterranean, heterothermic rodent distinguished by its unusual biology and striking appearance. Its exceptional physiological traits have made it a prominent model organism. Reflecting its scientific impact, *Science* named the naked mole-rat ‘Vertebrate of the Year’ in 2013. (courtesy of Bioparc Valencia, Spain).

### Experimental animals

Mole-rats are not commercially available and are classified by the International Union for the Conservation of Nature under the LC category (common or abundant species with no current risk of extinction). Postmortem brain samples used in this study were obtained from families maintained at the animal vivarium of the University of South Bohemia (Czech Republic) and from zoo-housed animals (Bioparc Valencia, Spain). Specimens ranged in age from newborn (e.g., 0 years old) to 13 years, which represents approximately mid-life for these species (Table 1). Most samples correspond to female individuals; only two males were included (one *H. glaber* and one *F. mechowii*). Experimental procedures involving animals at the University of South Bohemia complied fully with European legislation on animal care (EU Council Directive 2010/63/UE) and were approved by the local Ethical Committee for Animal Experimentation of the University of South Bohemia (ref. MZE-53257/2024-13143). Samples obtained from zoo-housed animals correspond to individuals that died from natural causes or from aggression or fighting among colony members. We confirm that we have read the Journal’s position on issues involved in ethical publication and affirm that this work is consistent with those guidelines.

**Table 1:** Experimental specimens.

| Species | Number | Sex | Age (years) |
|---|---|---|---|
| *H. glaber* | 3 | F | 0 |
| *H. glaber* | 3 | F | 2 |
| *H. glaber* | 4 | F | 4 |
| *H. glaber* | 2 | 1F + 1M | 7 |
| *H. glaber* | 1 | F | 11 |
| *H. glaber* | 1 | F | 13 |
| *F. mechowii* | 1 | M | 2 |
| *F. mechowii* | 1 | F | 7 |
| *F. mechowii* | 1 | F | 11 |

### Necropsy and tissue processing

Terminal anesthesia was induced with an overdose of a mixture of ketamine and xylazine. Brains were then removed from the skull and post-fixed for two weeks in a buffered 4% paraformaldehyde solution. After fixation, brains were stored for an additional week in a cryoprotectant solution containing 20% glycerin and 2% dimethylsulphoxide in neutral PBS. Frozen coronal sections (40

μm-thick) were obtained using a sliding microtome and collected in 0.125 M PBS (pH 7.4) as 10 series of adjacent sections spanning the entire rostrocaudal extent of the brain. These series were used for (i) direct visualization of NMel; (ii) Masson-Fontana staining; and (iii) multiple immunofluorescent detection of phosphorylated alpha-synuclein, P62, and tyrosine hydroxylase (TH), combined with brightfield visualization of NMel. A complete list of primary and bridge antisera is available at the Zenodo repository (https://doi.org/10.5281/zenodo.22791203). Sections were mounted on slides, air-dried overnight, and coverslipped with Entellan.

## 3 Results

An age-dependent pigmentation of catecholaminergic cell groups in the brains of *H. glaber* and *F. mechowii* was observed in the SNpc, VTA, and LC. Initial traces of NMel, barely above the detection threshold, appeared after two years of age and progressively increased, eventually occupying most of the cytoplasm. Across all specimens examined, the LC consistently displayed the highest levels of pigmentation, followed by the SNpc and the VTA (Figure 2). The catecholaminergic identity of pigmented neurons was confirmed by detection of the enzyme tyrosine hydroxylase (Figure 3). Although NMel granules were very small in two-year-old animals, granules of varying sizes (e.g., small and large granules) were observed in all catecholaminergic nuclei at all time points beyond four years of age (Figure 4). This heterogeneity likely reflects either distinct stages of NMel maturation or biochemical differences among pigmented catecholaminergic neurons. Indeed, multiple dopaminergic neuronal endophenotypes have been described in ventral mesencephalic cell groups in both rodent and primate brains *(22,23)*. In parallel, tier-specific selective pigmentation of SNpc dopaminergic neurons has been reported in mice and macaques, in which the most vulnerable ventral-tier neurons exhibited the highest pigmentation levels following the viral-mediated tyrosinase overexpression *(9,10)*.

**Figure 2.**
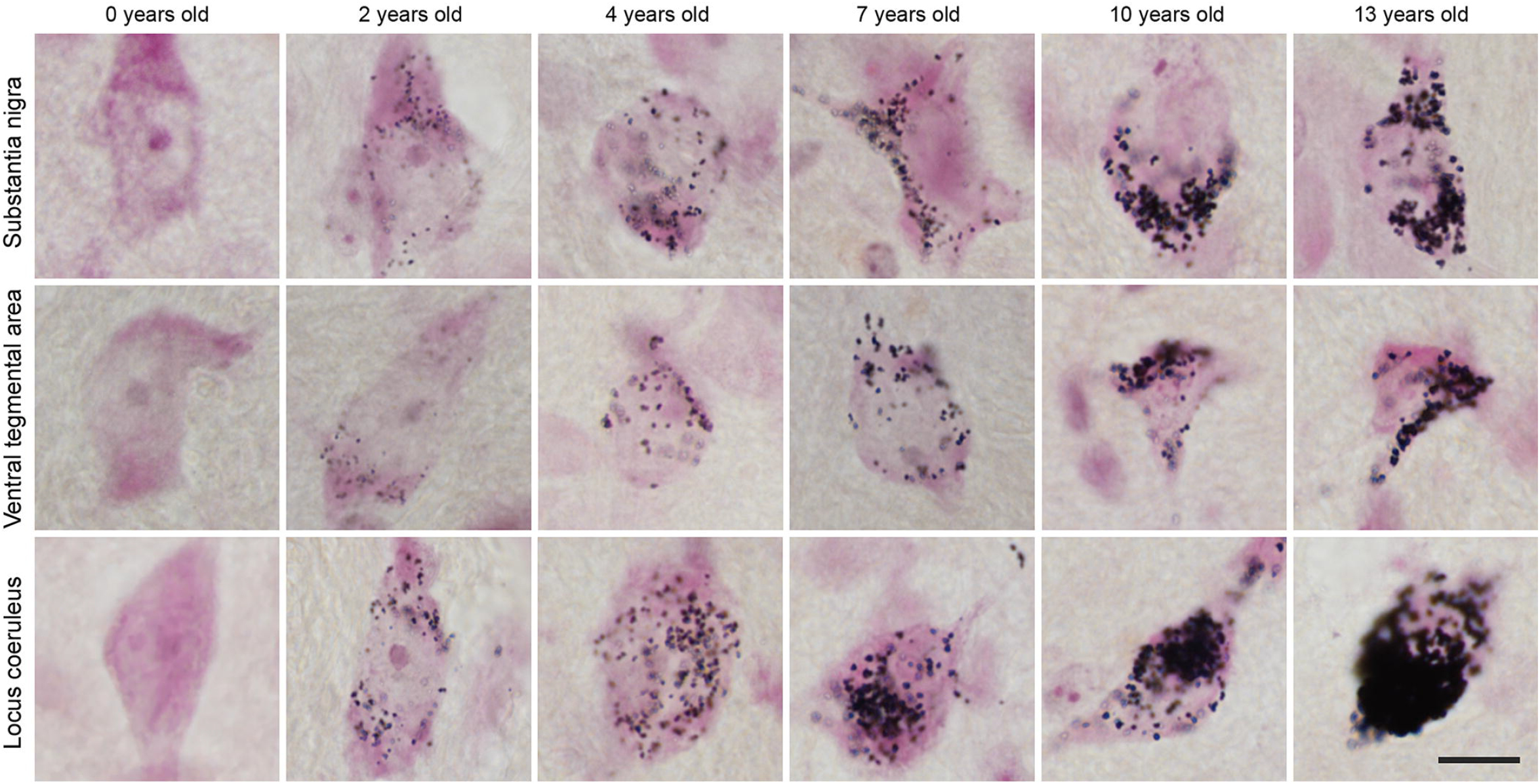
Time-dependent pigmentation of catecholaminergic centers in mole rats. High-magnification photomicrographs of brain sections stained with the Masson-Fontana histochemical method showing progressive, age-dependent pigmentation of dopaminergic neurons in the substantia nigra pars compacta (SNpc), ventral tegmental area (VTA), as well as noradrenergic neurons in the locus coeruleus (LC). At all examined ages, LC neurons exhibit the highest pigmentation levels. Scale bar, 10 μm in all panels.

**Figure 3.**
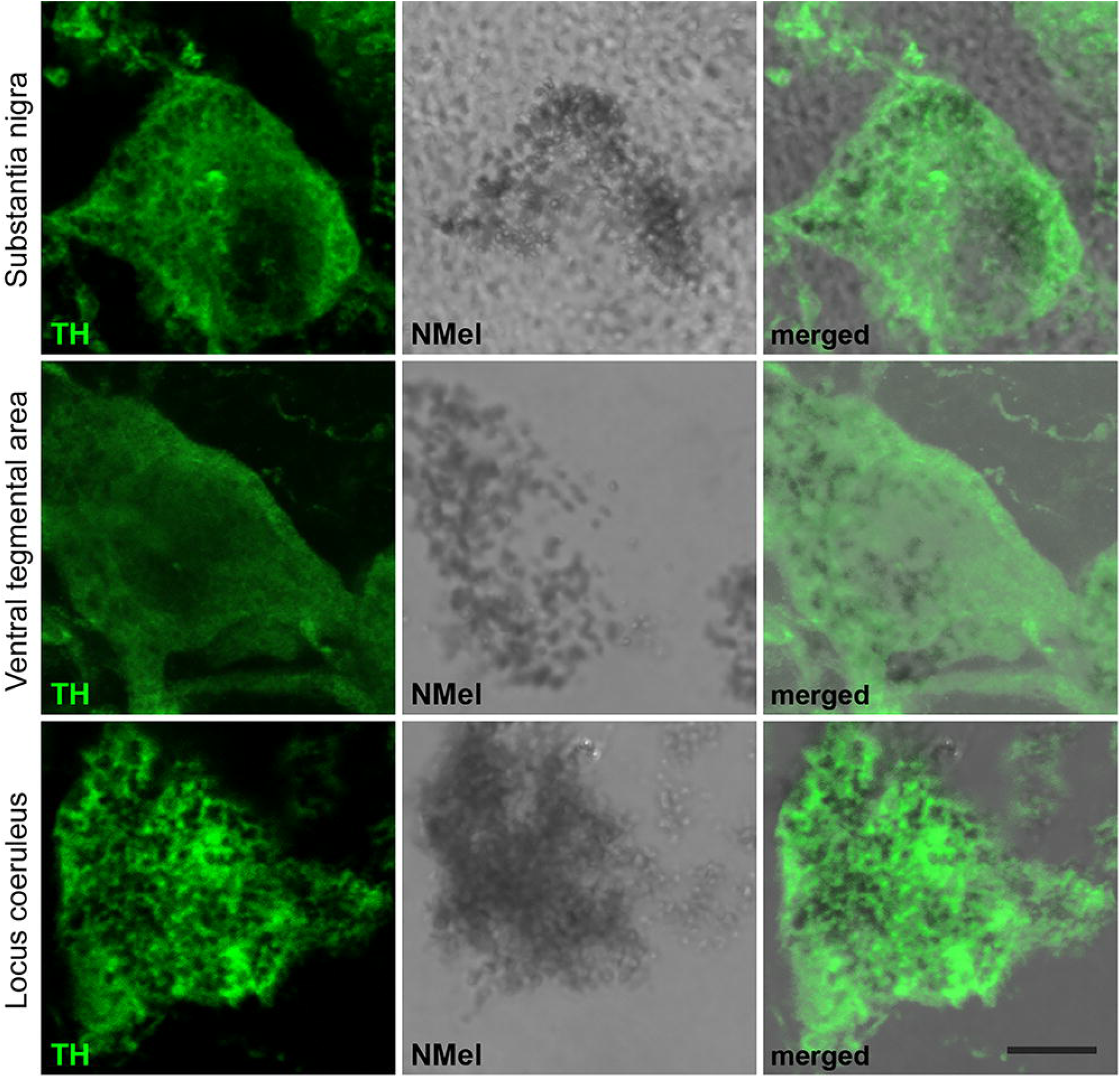
Catecholaminergic phenotype of pigmented neurons. Confocal high-magnification photomicrographs showing the catecholaminergic identity of pigmented neurons in the SNpc, VTA and LC, of a 10-year-old specimen. Only catecholaminergic neurons possess the molecular machinery required for NMel accumulation. Pigmented neurons are positive for the enzyme tyrosine hydroxylase (TH+; green channel). As in humans, the time-dependent accumulation of NMel in mole rats is physiological; accordingly, intracytoplasmic inclusions positive for canonical markers of Lewy body pathology -such as phosphorylated alpha-synuclein and P62-have never been observed. Scale bar, 5 μm in all panels.

**Figure 4.**
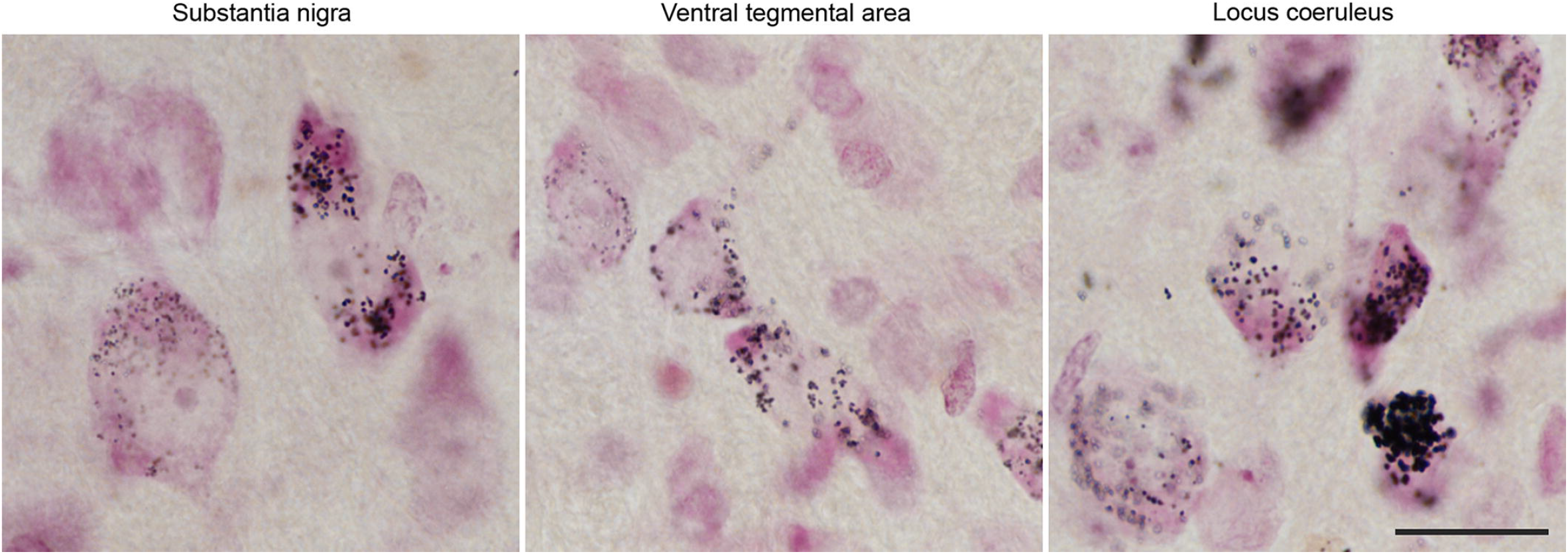
Varying sizes of neuromelanin granules. Both small and large NMel granules were observed in all inspected catecholaminergic centers in animals older than four years, likely reflecting distinct maturation stages of NMel accumulation and/or biochemical differences among catecholaminergic neurons. Scale bar, 30 μm in all panels.

## 4 Discussion

The question of why NMel synthesis is so slow, even in the presence of the required enzymatic machinery, remains unresolved. An enzymatic pathway involving tyrosinase has been proposed, given its established role in skin melanin synthesis from L-DOPA *(24)*, and viral-mediated tyrosinase overexpression has enabled the development of pigmented animal models of PD *(7-10)*. However, the presence of tyrosinase protein in the adult human SNpc remains controversial *(25-30)*, and tyrosine hydroxylase has likewise been proposed only as a potential contributor to NMel synthesis *(31)*. Crucially, despite the presence of both enzymatic and non-enzymatic pathways in rodent brains, catecholaminergic cell groups in mice and rats remain unpigmented. This suggests that NMel synthesis is governed by an intrinsically slow kinetic mechanism that these enzymes are unable to substantially accelerate *in vivo*. Consistent with this view, the non-enzymatic pathway, in which cytosolic dopamine auto-oxidizes to o-quinones, generating aminochrome, 5-S-cysteinyldopa and 5-5-cysteinyldopamine as precursors of the melanin components of NMel, may itself proceed too slowly to produce visible pigmentation within the lifespan of these animals.

Within the context of PD, evidence from three independent studies in which patients with PD received fetal midbrain cell transplants into the putamen supports the existence of a link among oxidized dopamine, NMel accumulation, and alpha-synuclein homeostasis *(32-34)*. Postmortem analyses revealed the unexpected presence of Lewy body-like inclusions positive for phosphorylated alpha-synuclein and ubiquitin within grafted neurons. These findings were interpreted as proof-of-concept for the prion-like spread of alpha-synuclein pathology from the host to the grafted tissue, and collectively represented a very important breakthrough in PD research. Although all these studies noted that transplanted neurons were moderately-to-heavily neuromelanized, the potential contribution of increased oxidative stress resulting from the conversion of cytosolic dopamine conversion into NMel, and its role in promoting alpha-synuclein aggregation, was largely overlooked. In this regard, it is worth noting that the age-dependent accumulation of NMel observed in mole-rats and reported here is physiological; accordingly, alpha-synuclein aggregates in the form of Lewy bodies were not observed.

In conclusion, this work addresses a deceptively simple yet unexplored question: why is NMel largely absent in rodents? Because NMel accumulates slowly over time, the relatively short lifespan of commonly used rodent species prevents the pigment from reaching detectable levels, even though their catecholaminergic neurons possess the molecular machinery required for its synthesis. By contrast, the exceptionally long longevity of mole-rats provides the temporal opportunity for NMel to accumulate to detectable levels. Thus, obtained findings indicate that mice and rats lack pigmentation not because they are incapable of synthesizing NMel, but probably because they do not live long enough for the pigment to accumulate to detectable levels. Indeed, although most commonly used laboratory rodents lack pigmentation in their brains, a few NMel-like granules have been detected by electron microscopy in rats aged more than two years *(35)*. Moreover, traces of NMel were found in mammals other than mice, rats and guinea pigs, such as carnivores, goats, adult horses and dolphins *(3, 36-39)*.

## 5 Article types

Brief Research Report.

Research topic *“Neuromelanin and Iron in Parkinson’s Disease: Mechanisms, Heterogeneity, Biomarkers, and Progression”*.

## 7 Author Contributions

Conceptualization & design: DR, JLL

Execution: CC, MMI, JC, AJR, GA, EL-R, DC, RS, JO

Analysis: CC, MMI, JC, GA, AJR, JH, DC, RS, JO, AJR

Funding acquisition: JLL Supervision: AJR, DR, JLL

Writing – original draft: CC, MMI, JH, JC, AJR

Writing – review & editing: AJR, DR, JLL

## 12 Data Availability Statement

Further information and requests for resources and reagents should be directed to and will be fulfilled by the corresponding author, Jose L. Lanciego. The raw data and key laboratory materials used and generated in this study are listed in a Key Resource Table alongside with their persistent identifiers are publicly available at the Zenodo repository (https://doi.org/10.5281/zenodo.22791203). A preprint version of the manuscript has been posted in a publicly accessible repository (https://www.biorxiv.org/content/10.64898/2026.10.13.759023v1).

